# Climate change velocity drives rapid evolution of foliar phenology in trailing edge populations

**DOI:** 10.64898/2026.04.27.719373

**Authors:** Shannon L.J. Bayliss, Ian M. Ware, Jennifer A. Schweitzer, Joseph K. Bailey

## Abstract

Here we tested the overarching hypothesis that climate change velocity drives rapid evolution in bud break phenology. With field and common garden studies, we used age cohorts within 17 populations of a foundation riparian tree species distributed across multiple strong environmental gradients in the western US. We provide evidence of contemporary evolution, as young trees in trailing-edge populations have evolved to break bud approximately six days earlier than old trees in these same populations. These populations experience greater water stress than populations at the core of the species’ distribution, and the magnitude of genetic divergence in bud-break phenology is related to the velocity of change in climate water deficit over the last 100 years. This relationship to climate change velocity did not exist in core populations, suggesting that, despite similar rates of change, a threshold of water deficit stress has yet to be surpassed in those regions. Overall, the interactive effects of old trees and trailing edge populations can provide useful insight into centuries of environmental history and each independently represent important benchmarks for understanding the context for contemporary environmental change.

## Introduction

Current projections suggest that climatic changes are outpacing the speed at which many plant species can shift extant populations^1,2,3^. Thus, populations found along the trailing edge (or the receding edge) of a shifting species distribution are predicted to retract as climates warm, ultimately persisting as climate relicts^4^. Two hypotheses have been proposed for how relict populations persist: 1) climate-driven selection resulting in adaptation or, 2) phenotypic plasticity in traits that are best suited for new climatic conditions. Evidence for contemporary evolution in response to changing climates in natural populations remains scarce and may be due, in part, to the misconception that evolutionary time scales are far slower than that of climatic change^5^. However, there must be capacity for contemporary evolution, alongside phenotypic plasticity, if *in situ* tree populations are to persist or adapt to the new climatic environments, they experience^6,7,8^. Species that occupy large geographical ranges can experience high abiotic and biotic environmental variation, resulting in differences in quantitative trait variation, population genetic differentiation, and adaptation to climate^9–12^. Understanding whether *contemporary evolution* is a significant factor in how tree species respond to modern climate change, how patterns of contemporary evolution vary geographically, and whether it is consistent across genetic lineages will improve how we understand and predict the magnitude of rapid evolutionary responses.

The strength and direction of climate change as a selective force varies geographically^13–16^. Individual climate gradients such as temperature and precipitation can diverge geographically and show discrepancies in their magnitude and direction of change over time^13,14,17^ due to land-surface processes and interactions. For example, the interaction of temperature and precipitation drive atmospheric processes like potential evapotranspiration (PET), and that the atmospheric demand for water from plants also varies geographically^18,19^. To better capture these complexities, it is useful to investigate multivariate climate variables, like PET or water budget deficit, and their associated local climate change velocity which describes the rate and magnitude of change at any point in space and time. This can help identify areas where organisms are exposed to rapidly changing climate^20^. The concept of climate change velocity,^21^ often applied in species range shift studies, also has high applicability to conservation-related research more broadly. Climate change velocity metrics are spatially explicit and provide a more comparable or standardized way to compare the degree of environmental change that populations experience across different landscapes, accounting for historical conditions and spatial heterogeneity reference 21. These spatial considerations are key when exploring how population-level genetic processes are occurring on landscapes with different historical conditions. Measuring local climate change velocity in a variable that considers both temperature and precipitation (i.e., PET, water budget deficit^22^, etc.) helps escape the issue of local divergences between the individual metrics of temperature and precipitation and allows the velocity metric to be interpretable as an organism’s exposure to actual climate change. However, the exposure, sensitivity, and vulnerability of species to the magnitude of climate change also depends on the species’ adaptive capacity (i.e., the ability of populations to intrinsically adjust to environmental changes, either through genetic changes, phenotypic plasticity and/or dispersal), especially for less mobile organisms like trees.

Areas of higher climate change velocity may be expected to force higher selective pressures on organisms, especially if changes to climate fall outside of the species range of thermal or drought tolerance^14^ and create novel climatic conditions. In this study, our focus is not on linking trait changes to specific environmental drivers directly, instead, we use climate change velocity to explore whether rates of trait change and rates of climate change are related across the landscape.

Building upon previous research demonstrating climate-driven evolution of spring bud break phenology and tree productivity at a landscape scale^12^, we sought to test the overarching hypothesis that climate change velocity drives rapid evolution in foliar bud break phenology. To test this hypothesis, first, we grouped 17 populations of foundational, riparian forest tree species, *Populus angustifolia* into “trailing-edge” (*n*=6) and “core” (*n*=11) groups based upon geographic barriers and isolation^2,3^. Second, using diameter at breast height (DBH) from the field, we established a tree size-age relationship^23^ to examine how phenology and productivity estimates measured in an experimental greenhouse common garden may have evolved through time and vary among populations geographically. Finally, to estimate climate change velocity, we extracted climatic water deficit and actual evapotranspiration values (ClimateNA^22^) for the period 1916-2005 because our “old” cohort of trees established at the earlier end of this period (~1940) and our “young” cohort of trees established closer to the end of this period (no earlier than 1987). Taking this integrative, biogeographic approach combining trailing-edge and core-populations with age cohorts we found: 1) evidence of contemporary evolution advancing spring phenology by an average of ~6 days in young trees from trailing-edge but not core populations; 2) reduced genetic variation in bud-break phenology in trailing-edge populations; 3) patterns of population genetic divergence and reduction in genetic variation consistent with a model of natural selection; 4) geographic variation in climate change velocities is related to atmospheric water deficit in core and trailing edge populations; and 5) ***the rate of evolutionary change in bud-break phenology is related to the rate of climate change in atmospheric hydrologic budgets over the last 100 years***.

## Results

To test for patterns of genetic divergence among age cohorts and range positions, we examined same-age cuttings taken from trees of varying ages in the field and planted in a common garden, When we characterized tree size in the field and bud break phenology and traits related to growth in the greenhouse common garden, we found bud break phenology in the common garden was related to diameter at breast height (DBH), an age corollary in the field, but only in trailing edge populations (Table S1 and S2, Figure S1). These results suggest that phenology is evolving in the trailing-edge, but not in core populations.

To test the hypothesis that young trees were diverging from old trees in trailing edge populations, in a common garden, we grouped trees from core and trailing edge populations into *young* (i.e., <25.4 years old) and *old* (i.e., >72.6 years old) cohorts. Tree age was estimated from field diameter at breast height using a growth-age relationship of 0.5-1.0 cm/yr,^23^). Consistent with world-wide observations of advancing spring phenology, we show rapid evolution of bud-break phenology and shoot length in young relative to old trees in trailing edge populations (**Figure 1**). We show break bud on average occurs six days earlier than old trees in trailing edge populations (χ^2^_(1,4)_ =6.68, p=0.03, *n*=124) and was earlier than young (11 days) and old (8 days) trees in core populations (Full model results in Table S3 and S4). Importantly, there was no quantitative difference in bud break among young and old trees in core populations (χ^2^ _(1,9)_=3.079, p=0.214, *n*=139). Similarly, cuttings derived from young trees have 37% longer new shoots than cuttings derived from old trees in trailing edge populations (χ^2^_(1,4)_ =6.154, p=0.046, *n*=135) and longer shoots than young (69%) and old (75%) trees in core populations (Full model results in Table S3 and S4, Mean trait estimates in Table S5). No differences were detected in shoot length among young and old trees in core populations (χ^2^_(1,4)_ =1.428, p=0.48, *n*=330). Finally, consistent with patterns of genetic divergence in bud-break phenology, we found reductions in the coefficient of genetic variation (CV_g_) for bud-break phenology in trailing edge populations relative to core populations suggesting a pattern of directional selection (χ^2^_(1,16)_ =10.64, p=0.0011, *n*=17; Figure 1C). CV_g_ estimates were transformed to standardized z-scores for analyses.

**Figure 1.**
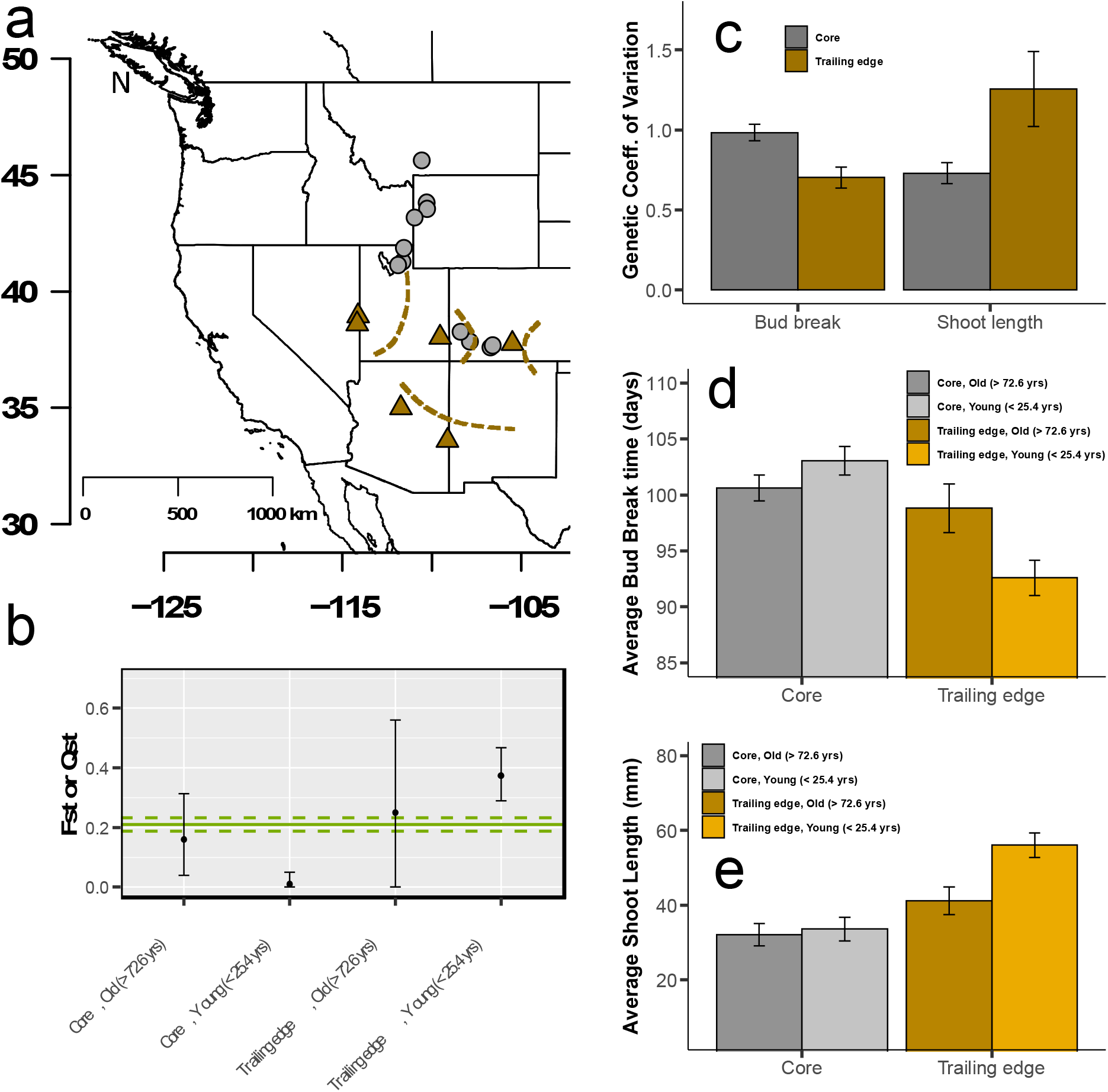
Contemporary evolution of bud break across tree populations. **Panel a** shows the distribution of core and edge populations along with geographic barriers represented by dotted lines; **Panel b** shows contemporary evolution of bud break, with the solid green line representing mean, global F_ST_ value for the *Populus angustifolia* populations observed. Dashed green lines represent 95% confidence intervals around the global F_ST_ estimate. Individual points represent mean Q_ST_ estimates for age cohorts (young and old) for both core and trailing edge populations. Error bars represent 95% confidence limits around the sample Q_ST_ means (Table S3. Global *F*_*ST*_ estimate generated using molecular data from 279 genotypes. **Panel c** shows the genetic coefficient of variation for population bud break and shoot length. Error bars represent ± one standard error of the mean. **Panels d & e** compare phenotypic trait variation among young and old trees across core and edge populations. Error bars in these panels represent ± 1 standard error of the mean.

To determine if detected genetic divergence in bud break phenology among age cohorts and range positions was driven by random genetic drift, natural selection, or a combination of both, we compared patterns of molecular evolution (*F*_*ST*_) to quantitative trait differentiation (*Q*_*ST*_). Population genetic differentiation (*F*_*ST*_), calculated from microsatellite data, was compared to quantitative genetic variation (*Q*_*ST*_) in foliar phenology within core and trailing edge populations and by age class^24–26^ (Figure 1B). Based on GenoDive results, there is strong overall molecular differentiation among all populations across the landscape (*F*_*ST*_ = 0.21, see Table S6). When we examine the older trees, regardless of whether they are found in core or trailing edge habitats, *F*_*ST*_ = *Q*_*ST*_, indicating that natural selection has not been a driving force affecting genetic variation in bud-break phenology for the older cohort. However, Qst<Fst for young trees from core populations and *Q*_*ST*_ > *F*_*ST*_ for young trees in trailing edge populations. These results support the hypothesis that stabilizing selection is acting on bud-break phenology in young trees in core populations. These results also support the hypothesis that directional selection is acting on young trees in trailing edge populations for earlier bud-break phenology.

In support of the hypothesis that patterns of rapid contemporary evolution are driven by climate change velocity and range position, we show that trees in trailing edge and core populations respond uniquely to the velocity of atmospheric water deficit (Table S7). Edge populations diverged in their phenological strategy the most in areas where the velocity of atmospheric water deficit was highest over the period of 1916-2005 (**Figure 2** and Table S8, scaled slope estimate +/−standard error = 1.05 +/−0.47, p=0.089). In other words, where drought conditions became the furthest from normal the fastest, evolution of bud-break phenology trended to occur the most. This pattern did not hold in the core population (Table S8, scaled slope estimate +/−standard error = −0.50 +/−0.32, p=0.16), though the climate velocity was highest overall in those regions (Figure 2A). Actual conditions in both the populations explain these patterns; the climate water deficit is much higher, and thus more stressful, in edge populations than in core populations (Figure 2B). In future climates, we may expect these patterns to continue in edge populations as our ecological niche models suggest that suitable niche space (i.e., niche breadth) will decrease over time for edge populations more than for core populations, regardless of emissions scenario (Figure 2D; Table S9).

**Figure 2.**
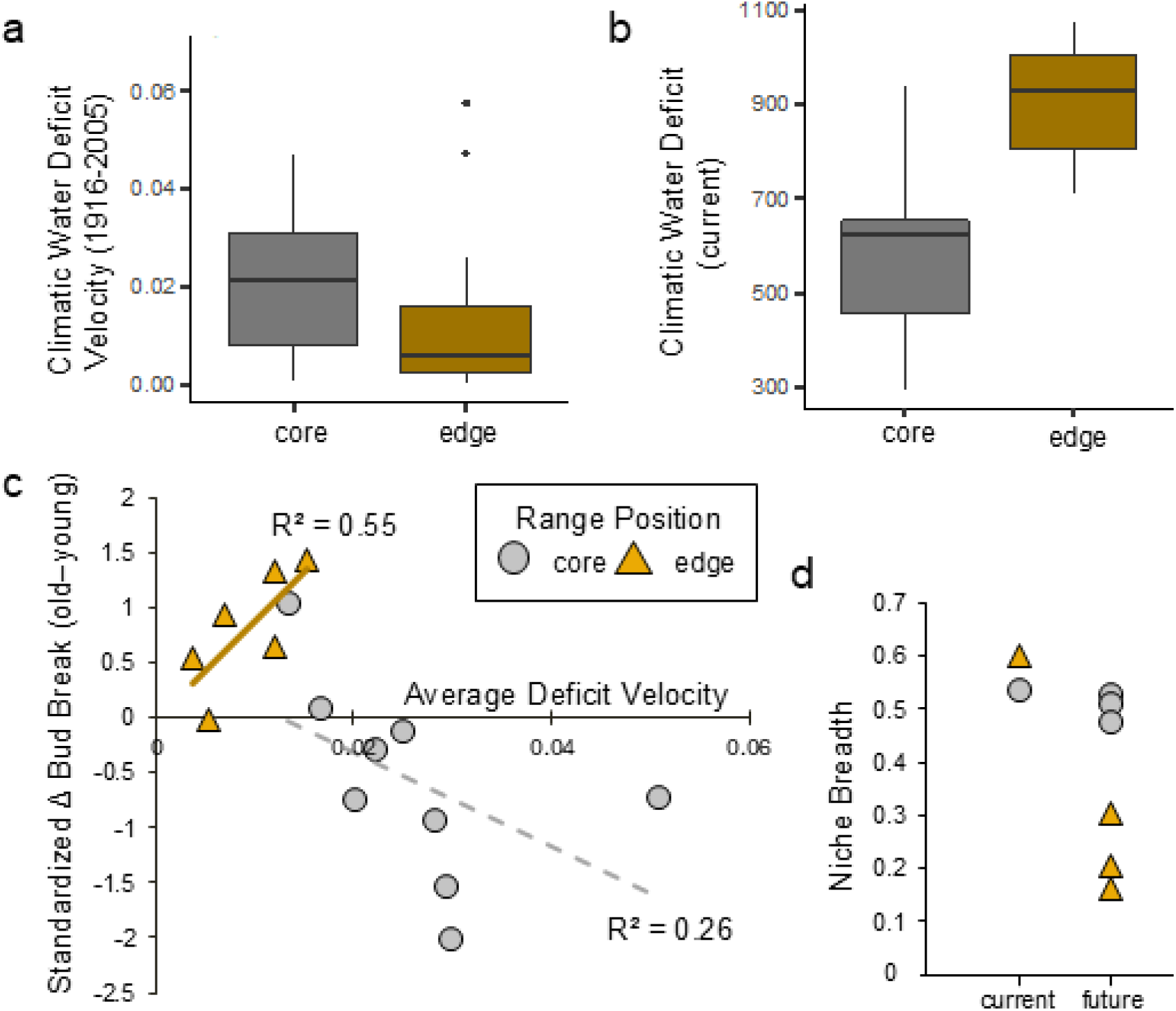
Climate driving contemporary evolution of bud break. **Panel a** represents climate deficit velocity from 1916-2005 across core and edge populations; **Panel b** shows current climate deficit across core and edge populations; **Panel c** represents the relationship between the average climate velocity represented in panel a with the degree of trait change between old and young trees in core and edge populations; **Panel d** represents predictions of future niche breadth across three global change models for core and edge populations.

## Discussion

Here we present evidence that geographic variation in the velocity of climate change is related to rapid evolution of phenology. Climate water deficit represents the difference between potential and actual evapotranspiration (PET and AET^27^) and is used as an indicator of drought^22^. Over at least the last 100 years, spring phenology has advanced by about six days in young trees from trailing edge populations. In addition to mean shifts in the timing of phenology, genetic variation in bud break phenology is reduced compared to core populations of the species. Both patterns of divergence and reduction in genetic variation are consistent with a model of rapid evolution occurring in trailing edge populations. We also find no difference between young and old trees from core populations, suggesting that we are not just capturing developmental differences between age cohorts. Over the same time period, the rate of change in climate water deficit, or the atmospheric demand for water from plants, has been geographically variable but absolute deficit values are highest in trailing edge populations. We note that climate change velocity calculated in mountainous regions with complex terrains can under-estimate climate change depending on whether a species’ minimizes exposure to different climates or travel distance^28^, though *Populus angustifolia* primarily disperses by wind and water, occurring along highly connected mountainous riparian corridors. These results are important because they show that there is enough genetic variation within and among trailing edge populations to respond to ongoing strong selective gradients associated with climate. Whether these populations continue to persist is still uncertain^4^.

We predict that the effects of climate change velocity on rapid evolution in bud-break seen over the last 100 years will continue over the next 100 years. Using ecological niche models, we show that niche breadth – the available suitable environmental space for the organism – is predicted to decrease in the edge populations in all three severity levels of future global change model predictions for the years 2071-2100 while core population lose niche breadth to a much lesser extent. This continued environmental stress is problematic for trailing edge populations that already have reduced genetic variation in bud-break phenology. These results are consistent with Bayliss et al. 2022^29^ that used a novel approach to model the spatial distribution of genetic variation across the landscape now and in the future. In that work, models predicted all populations would show earlier bud break phenology, which is consistent with worldwide observed trends in temperate regions, but that some populations will continue to lose genetic trait variation and undergo directional selection while others may experience stabilizing selection; patterns generally aligning with trailing edge and core populations.

Because the rate of evolution in bud-break phenology is related to the rate of change in climate water deficit hydrologic budgets over the last 100 years, there is potential for significant geographic variation in ecosystem-level climate feedback driven by differences in genetic variation. This is particularly true if there are genetic differences in traits that regulate or are correlated with water use^19^. This hypothesis is consistent with previous research suggesting that populations of *P. angustifolia* from hot and dry locations had stronger plant atmosphere feedbacks than populations from cool and wet localities^19^. Further, because of geographic variation in the atmospheric demand for water, populations with weak plant-atmosphere feedbacks (e.g., core populations) are likely to be under strong selection and represent zones of rapid evolution^27^ that may have large but under-appreciated impacts on atmospheric hydrologic cycles.

### Conclusions

We find strong support for the overarching hypothesis that climate change velocity in water deficit hydrologic budgets drives rapid evolution in bud break phenology. Comparing age cohorts within and among extant populations across the distribution of a focal species provides valuable insight into how important plant phenotypes are evolving on contemporary time scales to environmental pressures. The climate driven loss of genetic variation in a key ecological trait like bud break phenology that we show is significant. Bud-break phenology has implications for growing season length, biomass production, and resource availability for associated communities. Our results extend our understanding of eco-evolutionary dynamics of a long-lived foundation tree species that has direct genetically-based consequences for patterns of biodiversity and ecosystem functioning - including the conditioning of soil microbiomes^28–30^, insect communities, trophic interactions with insectivorous birds^31,32^, soil carbon and nitrogen pools and cycles^12,30,35,36^, and landscape scale plant-climate feedbacks through genetically based variation in hydrologic cycles^18^. Lastly, large, old trees are declining globally^37^ and are an amazing resource for understanding population genetic change and adaptive capacity to environmental change^38^. Old trees span centuries of environmental variation and should be a global conservation priority.

## Methods

### Study species, Site Selection, and Field Survey

*Populus angustifolia* James (Salicaceae) is a dominant tree species distributed throughout high elevation riparian zones (900 to 2500 m) along the Rocky Mountains from southern Alberta, through the intermountain U.S., and into northern Mexico^39^. Contemporary migration and population expansion is believed to be present in northern and central *P. angustifolia* populations, with increasing geographic isolation and population reduction in southern populations^40^, which supports the biogeographic models of Hampe and Petit^9^ and Woolbright et al.^4^. Further, individual river basins function as distinct genetic populations since gene flow among geographically separate forest stands is greatly reduced by geographic barriers, climatic factors, and the obligate riparian nature of *P. angustifolia* distributions^40^. Larger geographic features along the southern end of the geographic distribution further separate several populations from other major riparian corridors that could allow for gene flow among populations. The Great Basin spans the western and northwestern edge of the range, the San Juan Mountains separate the Colorado Plateau populations, and the Mogollon Rim separates the Arizona populations from both the Great Basin and Colorado Plateau (see dotted lines in Figure 1). In May 2012, seventeen distinct *P. angustifolia* river basins were surveyed collectively from three different genetic provenances (Arizona, Eastern, and Northern/Wasatch Clusters^39^)) across a gradient of ~1700 km latitude from southeastern Arizona to south central Montana. Collection sites occur across a large climatic range of approximately 10.4 degrees in mean annual temperature (−0.1°C to 10.3 °C) and 67.8 cm in annual precipitation (21.7 cm to 89.5 cm). See Ware et al.^12^ for more information regarding field observations, climatic data parameters, and data extraction).

To capture the range of genetic variation that occurred in each river basin, we identified and sampled from 3-5 collection sites: the highest and lowest elevation site with *P. angustifolia* trees and 1-3 intermediate locations within each river riparian area. Size of individual trees was recorded by measuring the diameter at breast height (DBH). Since *P. angustifolia* can propagate sexually and vegetatively, we sampled trees that were at least 30 m apart to avoid measuring and collecting genetic clones. Morphological differences in size, phenology, and growth form were also used to differentiate clones in the field. Tree age was determined by grouping DBH measurements into young and old cohorts. Young trees were determined as having a DBH in quartile 1 (i.e., 25% quartile and below), and old trees possess a DBH in quartile 3 (i.e., 75% quartile and above). To non-destructively determine the approximate age of each tree in the study, one year of growth was estimated to be ~ 0.50 cm increments in diameter. This was estimated from a growth-age relationship determined for *P. angustifolia* in the field^23^. We appreciate the potential for high variability in age-DBH relationships^41^, thus the binning of age cohorts rather than relying on potentially inaccurate estimates of ages for individual trees. Young trees were estimated to have established less than ~25.4 years ago (<12.7 cm in diameter), and old trees were estimated to have established more than ~72.6 years ago (>36.3 cm diameter). Field-collected leaf samples were oven dried (@ 70°C for 48 h) and ground into fine powder with a SPEX SamplePrep 8000D Dual Mixer/Mill (SPEX SamplePrep, Metuchen, NJ, USA). For 279 of the individual trees sampled, powdered sub-samples of leaf tissue were used to extract genomic DNA (gDNA) to determine genotypes and quantify population-level genetic divergence (*F*_*ST*_). (Extraction techniques and microsatellite information described in depth in^12^).

### Common garden and genetic differentiation

Common garden experiments are a powerful tool to explore the cause of phenotypic differences among populations. Clonal replicates of 582 field-surveyed genotypes were established in a greenhouse common garden in 2012 to minimize environmental effects and examine the genetic basis of functional plant phenotypes^42,43^. If phenotypic differences persist in a common garden environment, then there is evidence for genetic differentiation between the observed populations at loci that affect the trait(s) of interest.

Saplings grew for two years (quadrupling in size) in ambient light conditions with weekly water and monthly fertilizer during the growing season for maintenance (a water-soluble, balanced 20-20-20 of N, P, K). Ultra-Pure Oil Horticultural Miticide/Insecticide/Fungicide treatments were applied before bud break, after leaf senescence, and as needed to control fungal and pest outbreaks. The greenhouse common garden was located at the University of Tennessee in a climate-controlled glass greenhouse programmed to mimic seasonal changes in temperature. Two to four replicate saplings were selected at random from each surviving genotype to measure multiple plant traits associated with plant growth. In 2014, foliar bud break phenology (*n*=1,032 total plants) was measured every 48 h until all trees had flushed by recording bud break as the ordinal day when new leaves unfurl during spring emergence and represents the onset and ultimately the total accumulation of annual aboveground biomass production^44,44^. Additionally, before leaf senescence, shoot length (mm) was measured on the current year’s growth of the longest stem to provide an estimate of annual growth.

### Statistical Analyses

*Phenology and DBH correlations*. Exploratory trait correlations were performed to understand if phenology and tree diameter relationships changed due to geographic range position. We used a linear mixed-effects model with field-measured diameter at breast height (DBH, cm), range position (trailing-edge v. interior/core), and the interaction between DBH and range position included as fixed effects, and site (i.e., distinct field observational sites) included as a random effect. We included site to account for intraspecific and environmental variation within populations. Since DBH measurements were used to estimate age, these correlations explore geographic patterns in age-based genetic differences in phenology measure in a common garden.

#### Quantifying genetic variation, divergence, and differentiation for foliar phenology and annual growth

We calculated quantitative genetic variation within observed populations using plant clonal replicates in the common garden to estimate the possibility for selection. We calculated the genetic coefficient of variation (CV_G_) to estimate evolvability using the methods in^46^: CVG= 100 x V_G_/*x*, where V_G_ is genotypic variance and *x* is the population mean trait value. Genotypic variance (V_G_) of bud break phenology within each tree population was calculated using the var() function in base R. Calculating within-population genetic variation of phenotypes provides a conservative estimate to explore genetic clines in plant traits along landscape-level environmental gradients and are known to be good estimates of adaptive potential and evolvability^47^. CV_g_ estimates were transformed to standardized z-scores for regression analyses. Additionally, estimating within-population genetic variation can directly address hypotheses regarding patterns of genetic variation across a plant’s geographic distribution^4,9^.

To explore phenotypic variation among age cohorts, we estimated the mean phenotypes of saplings in the greenhouse common garden collected from young and old trees in natural populations. A mixed effects model was used to determine if bud break phenology and growth significantly differed in young and old trees across warm and cool habitats. In this model, range position (i.e., trailing edge and interior/core), age cohort (i.e., young and old), and the interaction of range position and age cohort were included as fixed effects; the population was included as a random effect. Tree population was included as a random effect to account for variation explained by genetic differences in phenotypes among populations. Once a significant interaction effect was detected, models were split to look for individual differences within core and trailing edge populations. Determining a phenotypic difference between young and old trees in trailing-edge populations, and no difference between age cohorts in interior populations, would provide evidence that tree populations are evolving to the modern climatic conditions on the landscape.

To estimate whether any observed quantitative trait differences were due to selection, among population genetic differentiation (*F*_*ST*_) was compared to quantitative genetic variation (*Q*_*ST*_) for foliar phenology and plant growth among young and old cohorts and among trailing-edge and interior populations. Using nine neutral microsatellite markers (Table S4), we determined population genetic differentiation (*F*_*ST*_) using GenoDive 2.0b27^48^. A global *F*_*ST*_ estimate was used for comparisons^49^. Comparing QST estimates to a global *F*_*ST*_ test for spatially divergent phenotypic selection against the range-wide neutral genetic differentiation. We also determined pairwise differentiation for all 17 sampled populations using AMOVA (10,000 permutations) in GenoDive 2.0b27. We grouped pairwise differentiation data by range position to determine neutral genetic differentiation within range position pairings (i.e., trailing-edge to trailing-edge, or interior to interior).

We determined quantitative trait differentiation (*Q*_*ST*_) among populations, using the same replicated genotypes from the common garden environment. *Q*_*ST*_ is a quantitative genetic analogue of *F*_*ST*_ that measures the amount of genetic variance among populations relative to the total genetic variance in a quantitative trait (rather than at a specific locus, as in the case of *F*_*ST*_). Following methods from a similar study in *Populus trichocarpa, Q*_ST_ and its 95% confidence intervals were estimated by resampling populations^48^. Individual *Q*_*ST*_ values were calculated using the following model, where 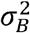 represents between-population variance and 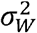 represents within-population (i.e., genotype-level) variance^24,25^:

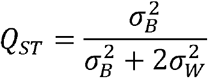

Following a commonly used technique, *F*_*ST*_-*Q*_*ST*_ comparative studies are used as an exploratory tool to determine if specific quantitative traits are under selection or are no different than if acted upon by genetic drift alone, or random, chance shift in phenotype. When *F*_*ST*_=*Q*_*ST*_, trait divergence among populations could be attributed to genetic drift alone, suggesting difference among populations are no different than random genetic change. If *F*_*ST*_>*Q*_*ST*_, trait divergence exceeds neutral expectations and is likely a product of directional or divergent selection, in which an extreme phenotype is favored over others. Further, if *F*_*ST*_<*Q*_*ST*_, trait divergence is less than expected by genetic drift alone and is suggestive of stabilizing or uniform selection across populations, in which genetic variation decreases and the population mean stabilizes around a particular phenotype^24–26^. We predict high levels of population genetic differentiation (*F*_*ST*_) based on previous work within *Populus* using some of the same microsatellites (see Evans et al.^39^ and citations within).

Determining quantifiable differences in *F*_*ST*_ - *Q*_*ST*_ estimates will allow for conclusions of detectable evolution regarding selection on phenotypic traits. If surrounding 95% confidence intervals associated with our estimates of differentiation do not overlap, we assume significant difference between observed estimates of population genetic differentiation (*F*_*ST*_) and quantitative trait differentiation (*Q*_*ST*_), providing phenotypic evidence of selection. We predict *F*_*ST*_-*Q*_*ST*_ comparisons will help identify phenotypic traits under selection in trailing-edge populations, providing evidence for climate-driven selection in spring phenology.

#### Quantifying the relationship between climate change velocity and phenological trait change in the trailing edge and core populations

We used climate change velocity metrics from ClimateNA which describe the exposure of organisms to climate change on the landscape (ClimateNA,^22^. ClimateNA provides past climate change velocity information for climatic water deficit and actual evapotranspiration. We specifically extracted values from the period 1916-2005 because our “old” cohort of trees established at the earlier end of this period (~1940, >72.6 years from 2012 collection) and our “young” cohort of trees established closer to the end of this period (no earlier than 1987, <25.4 years from 2012 collection). With the expectation that the faster the velocity of change, the greater the difference in traits between old and young trees, we ran linear models with the absolute value of phenological trait change between old and young trees as the response, and with climate velocity and population (core or edge) as predictors.

*To predict how core and edge population climate niche breadth will change in the future*, we built ecological niche models for both populations and projected them into 3 future scenarios to predict how the breadth of environmental conditions available to each population will change over time. To build the ENMs, we split occurrence data into two populations (core and edge). We checked for duplicate records then ensured no pseudo-replication in models by maintaining only one occurrence record for each cell of environmental raster data used to build models (resolution = 0.04166). We created geographic background extents for training the two models by estimating minimum convex polygons around each set of occurrence points using the function “mcp” in the R package adehabitatHR^50^. We added a buffer around the edges of the minimum convex polygons, so no occurrence point occurred on the edge of the geographic training extent using the “buffer” function in the R package “raster”^51^. Bioclimatic variables from 1961-1990 from the AdaptWest Project^52^ [v.7.21] were used to build models. The AdaptWest Project uses data from PRISM and WorldClim to develop informational resources to plan for climate adaptation in North America^52–54^. We ran initial models using Maxent (Version 3.4.1)^55^ with variables that had previously been determined to be of importance in describing the distribution of this species^27^. For each population, we removed any variables correlated above a threshold of 0.7 and ran five model replicates from which we maintained variables that cumulatively contributed 90-95% to the models to build final models. Final models were built with five-fold cross-validation and projected into future ensemble predictions (13 global change models) for the years 2071-2100 (ssp 245, 370, and 585 - increasing in severity). Average test AUC plus/minus standard deviation for edge population models was 0.83 +/−0.04 and for the core population models was 0.82 +/−0.02 indicating good model performance^56^. We calculated niche breadth for current conditions and future conditions with ENMTools^57^. All analyses were performed in R^58^, unless otherwise noted.

## Acknowledgments

We would like to extend gratitude to Michael Van Nuland, Phil Patterson, Ken McFarland, Jeff Martin, Alex Neild, Dailee Metts, Kassie Hollabaugh, Kelsey Greiff, Parker Wilson, Kaleb Menzel, Dylan Johnson, Blake Scalf, Katie Baer, and Alex Gifford for lab and greenhouse support. Funding support for IMW, SLJB, JAS and JKB was from the University of Tennessee, Knoxville.

## Author contributions

IMW, SLJB, and JKB conceptualized the study. IMW performed the field work and data collection. IMW and SLJB performed the statistical analysis and drafted the manuscript. All authors discussed results and equally contributed to subsequent drafts of the manuscript.

## Data availability

Observational and experimental data generated during and/or analyzed during the current study can be found as Supplementary Data 1-4.

## Code availability

All code was performed in R (version 4.4.0). Code generated during this study are available from the corresponding authors upon reasonable request.

