## Supplemental Information for "Climate change velocity drives rapid evolution of foliar phenology in trailing edge populations"

**Running Title: Climate change velocity drives rapid evolution of phenology**

**Affiliations:**
[1] Department of Environmental Science, Policy, and Management, University of California, Berkeley, CA, USA.
[2] Funga PBC 1612 Canterbury Street, Austin, Texas, 78702, USA.
[3] Department of Ecology and Evolutionary Biology, University of Tennessee, Knoxville, TN, USA.

***Corresponding first authors:** Shannon L.J. Bayliss & Ian M. Ware

### Abstract
Here we tested the overarching hypothesis that climate change velocity drives rapid evolution in bud break phenology. With field and common garden studies, we used age cohorts within 17 populations of a foundation riparian tree species distributed across multiple strong environmental gradients in the western US. We provide evidence of contemporary evolution, as young trees in trailing-edge populations have evolved to break bud approximately six days earlier than old trees in these same populations. These populations experience greater water stress than populations at the core of the species' distribution, and the magnitude of genetic divergence in bud-break phenology is related to the velocity of change in climate water deficit over the last 100 years. This relationship to climate change velocity did not exist in core populations, suggesting that, despite similar rates of change, a threshold of water deficit stress has yet to be surpassed in those regions. Overall, the interactive effects of old trees and trailing edge populations can provide useful insight into centuries of environmental history and each independently represent important benchmarks for understanding the context for contemporary environmental change.

### Keywords
phenology, climate velocity, adaptation, biogeography, contemporary evolution, genetic variation – climate-evolution feedback*

### Supplemental Material

### Tables

**Table S1.** Full mixed effects model results for the response of bud break phenology (days, *n*=380) and shoot length (mm, *n*=404) to diameter at breast height (cm), range position, and the interaction between DBH and range position.

| ***Response: Bud break phenology*** | $X^2$ | ***DF*** | ***Pr(>$X^2$)*** |
|---|---|---|---|
| DBH | 0.021 | 1 | 0.886 |
| Range Position | 10.60 | 1 | **0.001** |
| DBH * Range Position | 4.157 | 1 | **0.0414** |
| ***Response: Annual Shoot Length*** | | | |
| DBH | 0.66 | 1 | 0.418 |
| Range Position | 41.09 | 1 | **<<0.00001** |
| DBH * Range Position | 5.20 | 1 | **0.0225** |

**Table S2.** Full mixed effects model results for Bud break phenology (days, n=404) and shoot length (mm, n=465) differences by range position, age cohort, and the interaction between range position and age cohort. Tree population was included as a random effect.

| ***Response: Bud break phenology*** | $X^2$ | ***DF*** | ***Pr(>$X^2$)*** |
|---|---|---|---|
| Range Position | 3.45 | 1 | 0.063 |
| Age Cohort | 1.25 | 2 | 0.5[illegible]3 |
| Range Position*Age Cohort | 6.62 | 2 | **0.036** |
| ***Response: Shoot length*** | | | |
| Range Position | 15.76 | 1 | **0.00007** |
| Age Cohort | 2.39 | 2 | 0.301 |
| Range Position*Age Cohort | 6.05 | 2 | **0.04** |

**Table S3.** $F_{ST}$ and $Q_{ST}$ estimates and associated confidence intervals old and young cohorts among range position groupings.

| Age | Range Position | $Q_{ST}$ or $F_{ST}$ | Upper CI | Lower CI |
|---|---|---|---|---|
| Trailing edge, Old (> 72.6 yrs) | Edge | 0.25 | 0.31 | 0.25 |
| Trailing edge, Young (< 25.4 yrs) | Edge | 0.3737 | 0.0933 | 0.084 |
| Core, Old (> 72.6 yrs) | Core | 0.16 | 0.154 | 0.12 |
| Core, Young (< 25.4 yrs) | Core | 0.0104 | 0.0396 | 0.0104 |
| Global $F_{ST}$ | -- | 0.21 | 0.2326 | 0.1874 |

**Table S4.** Full model results to determine the response of the change in phenology (Δ phenology) to climate deficit velocity, range position, and their interactive effects. All variables have been z-transformed prior to running the model.

| ***Response: Δ phenology*** | **Estimate + SE** | **DF** | **Pr (>\|t\|)** |
|---|---|---|---|
| Climate Deficit Velocity 1916-2005 | -0.50 +/- 0.27 | 1 | 0.089 |
| Range Position | 1.93 +/- 0.78 | 1 | 0.031 * |
| Climate Deficit Velocity * Range Position | 1.55 +/- 0.85 | 1 | 0.098 |
| Residual standard error: 0.68 on 11 degrees of freedom<br>Multiple R-squared: 0.634, Adjusted R-squared: 0.0.534<br>F-statistic: 6.355 on 3 and 11 DF, p-value: 0.0093 | | | |

**Table S5.** Individual linear models, split by range position, relating response change in phenology ($\Delta$ phenology) to predictor climate deficit velocity. All variables were z-transformed prior to running models.

| ***Model:*** Range Position | Climate Deficit Velocity 1916-2005 **Estimate + SE** | **DF** | **Pr (>\|t\|)** | **R2** |
|---|---|---|---|---|
| Core | -0.50 +/- 0.32 | 7 | 0.157 | 0.26 |
| Trailing Edge | 1.05 +/- 0.47 | 4 | 0.089 * | 0.55 |

**Table S6**. Climatic variables contributing to ecological niche models of the core vs. trailing edge populations**.**

| **Percent % Contribution of Variables to ENMs** | | |
|---|---|---|
| | **Core** | **Trailing Edge** |
| **cmi Hogg's climate moisture index (mm)** | 0.33 | 56.72 |
| **eref Hargreave's Reference Evaporation** | 25.86 | 0.07 |
| **map Mean annual precip. (mm)** | 19.68 | 15.63 |
| **mcmt Mean temp. of the coldest month (*C)** | 3.28 | 1.48 |
| **msp Mean summer (May-Sep) precip. (mm)** | 45.92 | 1.71 |
| **pas Precipitation as snow (mm)** | 2.16 | 20.85 |
| **td Continentality MCMT-MWMT (*C)** | 2.76 | 3.54 |

**Figures**

**Figure S1**

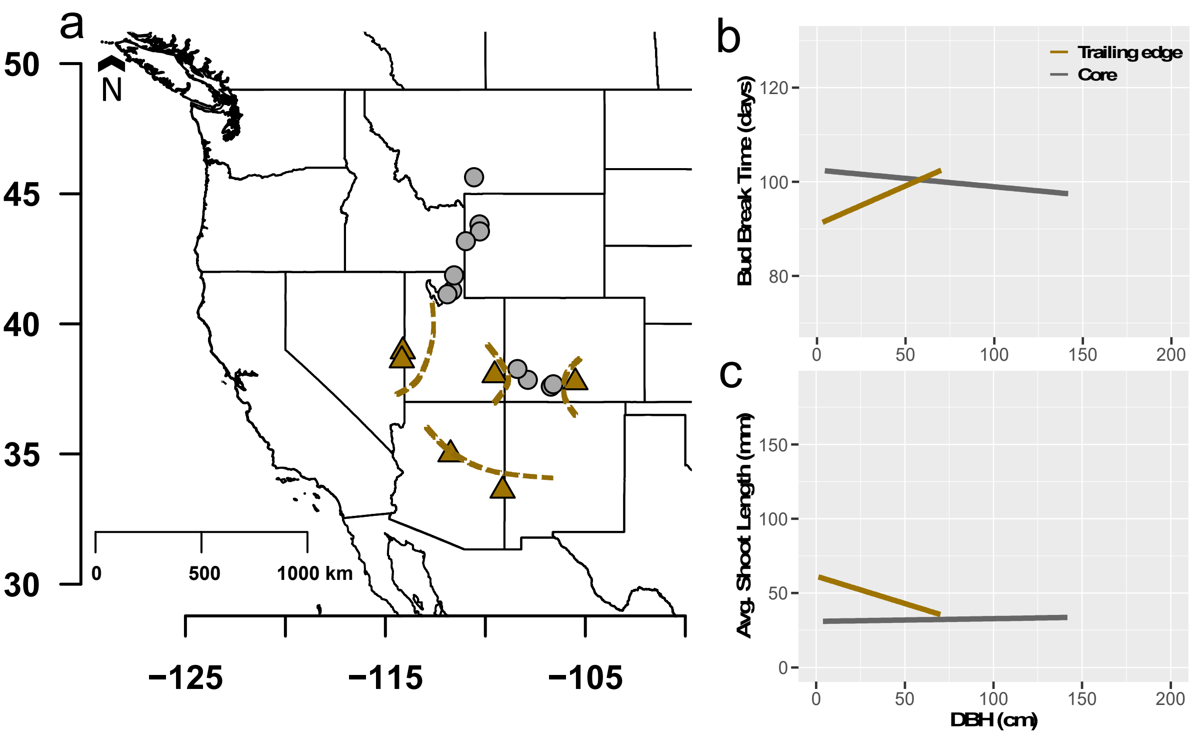

**Figure S2**

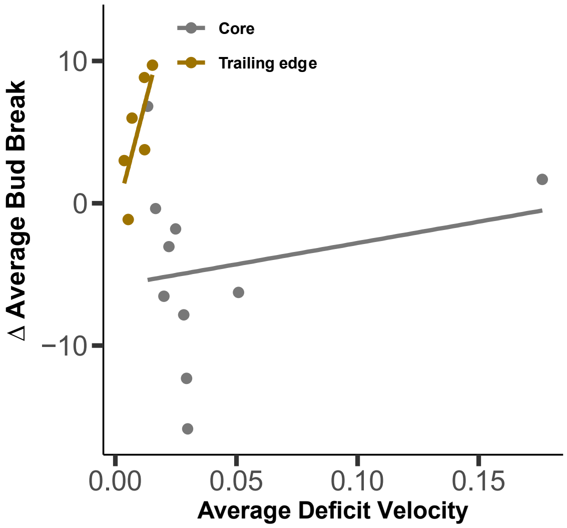
Core
Trailing edge
Δ Average Bud Break
10
0
−10
0.00
0.05
0.10
0.15
Average Deficit Velocity
